# Regional Striatal Cholinergic Involvement in Human Behavioural Flexibility

**DOI:** 10.1101/392233

**Authors:** Tiffany Bell, Michael Lindner, Angela Langdon, Paul Gerald Mullins, Anastasia Christakou

**Affiliations:** School of Psychology and Clinical Language Sciences, and Centre for Integrative Neuroscience and Neurodynamics, University of Reading, RG6 6AL, UK; Princeton Neuroscience Institute, Princeton University, NJ 08544, USA; School of Psychology, Bangor University, LL57 2DG UK

## Abstract

Animal studies have shown that the striatal cholinergic system plays a role in behavioural flexibility but, until recently, this system could not be studied in humans due to a lack of appropriate non-invasive techniques. Using proton magnetic resonance spectroscopy (^1^H-MRS) we recently showed that the concentration of dorsal striatal choline (an acetylcholine precursor) changes during reversal learning (a measure of behavioural flexibility) in humans. The aim of the present study was to examine whether regional average striatal choline was associated with reversal learning. 36 participants (mean age = 24.8, range = 18-32, 20 female) performed a probabilistic learning task with a reversal component. We measured choline at rest in both the dorsal and ventral striatum using ^1^H-MRS. Task performance was described using a simple reinforcement learning model that dissociates the contributions of positive and negative prediction errors to learning. Average levels of choline in the dorsal striatum were associated with performance during reversal, but not during initial learning. Specifically, lower levels of choline in the dorsal striatum were associated with a lower number of perseverative trials. Moreover, choline levels explained inter-individual variance in perseveration over and above that explained by learning from negative prediction errors. These findings suggest that the dorsal striatal cholinergic system plays an important role in behavioural flexibility, in line with evidence from the animal literature and our previous work in humans. Additionally, this work provides further support for the idea of measuring choline with ^1^H-MRS as a non-invasive way of studying human cholinergic neurochemistry.

**SIGNIFICANCE STATEMENT:** Behavioural flexibility is a crucial component of adaptation and survival. Evidence from the animal literature shows the striatal cholinergic system is fundamental to reversal learning, a key paradigm for studying behavioural flexibility, however, this system remains understudied in humans. Using proton magnetic resonance spectroscopy, we showed that choline levels at rest in the dorsal striatum are associated with performance specifically during reversal learning. These novel findings help to bridge the gap between animal and human studies by demonstrating the importance of cholinergic function in the dorsal striatum in human behavioural flexibility. Importantly, the methods described here can not only be applied to furthering our understanding of healthy human neurochemistry, but also to extending our understanding of cholinergic disorders.

## INTRODUCTION

Acetylcholine (ACh) plays an important role in adaptive behaviour, and has been implicated in disorders of cognitive flexibility, such as Parkinson’s disease (Tanimura et al., 2018; Zucca et al., 2018). Studies in rodents have repeatedly demonstrated that ACh transmission, determined by the activity and regulation of cholinergic interneurons in the dorsal striatum (DS), is involved in reversal learning and similar forms of behavioural flexibility (Ragozzino et al., 2002, 2009; Tzavos et al., 2004; McCool et al., 2008; Brown et al., 2010; Bradfield et al., 2013; Aoki et al., 2018; Okada et al., 2018). Further, ACh efflux has been shown to increase specifically during reversal learning (but not during initial learning), and this effect is specific to the dorsomedial striatum (with no changes in ACh levels in either the dorsolateral striatum or the ventral striatum) (Ragozzino et al., 2009). It is clear then that cholinergic activity in the DS plays an important role in reversal learning but, despite the importance of understanding this system, there remain important challenges in probing ACh function in humans due to a lack of appropriate non-invasive techniques.

Proton magnetic resonance spectroscopy (^1^H-MRS) is a non-invasive method for measuring brain metabolites *in vivo* (Puts and Edden, 2012). Although it cannot be used to study ACh directly due to its low concentration (Hoover et al., 1978), ^1^H-MRS can be used to measure levels of certain choline containing compounds (CCCs) involved in the ACh cycle, including choline (CHO). CHO is the product of ACh hydrolysis, and its uptake in cholinergic terminals is the rate-limiting step in ACh biosynthesis (Lockman and Allen, 2002). Using functional ^1^H-MRS we previously demonstrated task-driven changes in the concentration of CHO in the human DS during reversal learning (Bell et al., 2018). Although ^1^H-MRS studies typically model CCCs as a single peak due to their proximity on the spectrum, we showed that using this method may mask CHO-specific effects. Therefore, in the context of studying ACh function, it is necessary to separate the metabolites when measuring individual differences in CHO levels (Lindner et al., 2017; Bell et al., 2018).

Among the many open questions around this approach is the nature of the relationship between baseline levels of CHO availability and function-relevant ACh activity. Animal studies have shown that ACh synthesis is tightly coupled to CHO availability. For example, depletion of CHO has been shown to reduce ACh synthesis (Jope, 1979) and administration of CHO has been shown to increase it (Koshimura et al., 1990). Further, overexpression (Holmstrand et al., 2013) and under-expression (Parikh et al., 2013) of presynaptic CHO up-take transporters has been shown to increase and decrease ACh levels respectively. It is possible, therefore, that baseline CHO availability may modulate ACh activity, leading to effects on behavioural flexibility. In this study, we used ^1^H-MRS to test whether baseline levels of regional striatal CHO are related to individual differences in performance during a probabilistic reversal learning task. To do this, we obtained average measures of CHO from the dorsal and ventral regions of the striatum (DS and VS, respectively). Additionally, CHO levels from the cerebellum were used as a control to demonstrate specificity. In line with the animal literature and our previous findings in humans (Bell et al., 2018), we predicted that average levels of CHO in the dorsal, but not the ventral, striatum would be associated with performance during reversal, but not initial, learning.

## METHODS

### Participants

The study was approved by the University of Reading Research Ethics Committee. 36 volunteers (20 female) between the ages of 18.3 and 32.8 (mean = 24.8, SD = 3.5) were recruited by opportunity sampling. All participants were healthy, right handed non-smokers and written informed consent was taken prior to participation. Two participants were excluded from analyses due to a high proportion of missed responses (participant 14: 35% during initial learning and 39% during reversal learning; participant 31: 27% during initial learning, 54% during reversal learning).

### Behavioural Data

#### Learning Task

The task used was a probabilistic multi-alternative learning task previously described (Bell et al., 2018), and was programmed using MATLAB (2014a, The Mathworks, Inc., Natick, MA, United States) and Psychtoolbox (Brainard, 1997).

First, participants were presented with a fixation cross displayed in the centre of the visual display. Participants were then presented with four decks of cards. Each deck contained a mixture of winning and losing cards, corresponding respectively to a gain or loss of 50 points. The probability of getting a winning card differed for each deck (75%, 60%, 40%, and 25%) and the probabilities were randomly assigned across the four decks for each participant. Participants indicated their choice of deck using a computer keyboard. Outcomes were pseudo-randomised so that the assigned probability was true over every 20 times that deck was selected. Additionally, no more than 4 cards of the same result (win/loose) were presented consecutively in the 75% and 25% decks and no more than 3 cards of the same result in the 60% and 40% decks. A cumulative points total was displayed in the bottom right-hand corner throughout the session and in the centre of the visual display at the end of each trial (Figure 1). Participants were instructed that some decks may be better than others, they are free to switch between decks as often as they wish, and they should aim to win as many points as possible.

**Figure 1:**
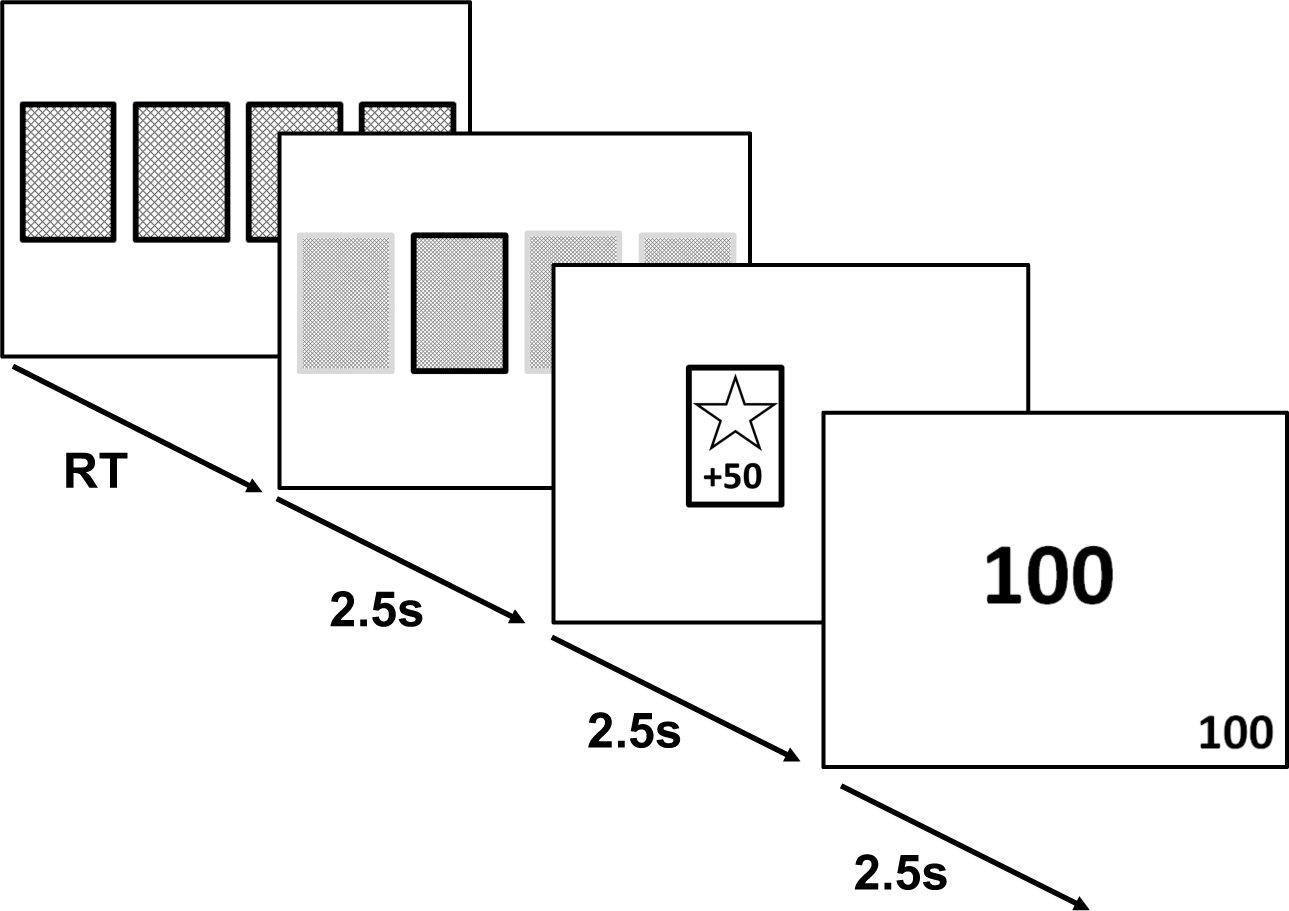
General outline of learning task trials. Participants were instructed to choose between four decks of cards. Each deck had a different probability of generating wins:losses (75:25, 60:40, 40:60, 25:75). Once the learning criterion had been reached, the deck probabilities were reversed so that high probability decks became low probability decks and vice versa. Participants were not informed of this in advance and were simply instructed to gain as many points as possible. Each screen was shown for 2.5s, RT = reaction time.

The learning criterion was set at selection of either of the two highest decks (60% or 75%) on at least 80% of the time over ten consecutive trials. Though the optimal strategy is to repeatedly choose the 75% deck, pilot testing revealed the participants were not always able to distinguish between the 75% and 60% decks. Therefore, as both decks generate an overall gain in points and choice of either deck could be considered a good strategy, both decks are included in the learning criterion.

The initial learning phase (round 1) was completed when either the learning criterion was reached, or the participant completed 100 trials. The deck probabilities were then reversed so that the high probability decks became low probability and vice versa. Participants were not informed of the reversal. The task ended either after the learning criterion was reached following the reversal (round 2), or after another 100 trials (Figure 2).

**Figure 2:**
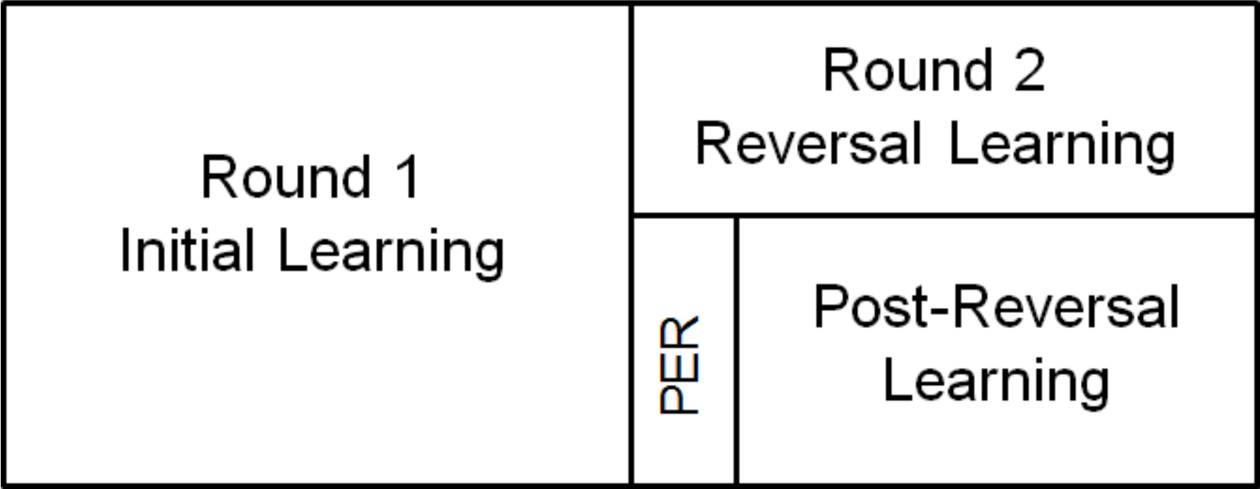
General overview of learning task structure. Participants completed the initial learning phase (round 1) by reaching the predefined accuracy criterion or after 100 trials. Upon completion of the initial learning phase, the deck probabilities were reversed. Participants then completed a reversal learning phase (round 2). For behavioural analysis, this was subdivided into perseverative trials (PER) and a post-reversal learning period. The number of perseverative trials was defined as the number of trials after reversal until the probability of selecting the previously favoured card reached chance level (0.25). The post-reversal learning period was the number of trials to reach criterion in round 2, minus the number of perseverative trials. The number of regressive errors was defined as the number of times the previously favoured deck was selected during the post-reversal learning period. The task ended once participants either reached the same accuracy criterion in round 2 or after 100 round 2 trials.

#### Impulsivity

Previous research has shown that trait levels of impulsivity can influence decision making (Bayard et al., 2011). Individuals with higher levels of impulsivity have been shown to demonstrate sub-optimal performance on decision making tasks, displaying a decreased ability to learn reward and punishment associations and implement these to make appropriate decisions. For instance, individuals with high levels of impulsivity were relatively impaired in adapting their behaviour during a reversal learning task (Franken, van Strien, Nijs, & Muris, 2008). Other tasks of cognitive flexibility have also been shown to be influenced by trait impulsivity levels (e.g. Müller, Langner, Cieslik, Rottschy, & Eickhoff, 2014). Therefore all participants completed the Barratt Impulsiveness Scale (BIS-11; Patton, Stanford, & Barratt, 1995) and their total score was used as a trait measure of impulsivity. This was included in the analysis to account for effects driven by individual differences in impulsivity.

#### Data Analysis

Participants were split into two groups based on performance. Those who learnt both rounds (i.e. reached criterion both during initial learning and after reversal) were classified as learners and those who did not learn both rounds were classified as non-learners.

Behaviour was analysed for learners only. Those who did not reach criterion during round 1 will not have realised there was a change in contingencies and will not have experienced a reversal, therefore their behaviour during the reversal stage is likely to be different to those who did experience the reversal. Additionally, because the task stops at 100 trials per round if the criterion is not met, there is a ceiling effect for those who did not reach criterion. Consequently, there will be a ceiling effect for those who did not reach criterion in both rounds, and for those who did not reach criterion in either round 1 or round 2 only. Therefore, participants who did not reach criteria in either one round or both rounds were excluded from behaviour analysis.

Performance was measured using the number of trials taken to reach criterion in round 1 (initial learning) and in round 2 (reversal learning). Round 2 was subdivided into perseverative trials and post-reversal learning (Figure 2). The number of perseverative trials was defined as the number of trials after reversal until the probability of selecting the previously favoured deck reached chance level (0.25), i.e. the number of trials taken to identify the reversal and switch behaviour. Post-reversal learning was defined as the number of trials taken to reach criterion in round 2, minus the number of perseverative trials, i.e. the number of trials to reach criterion after the reversal had been detected. In other words, post-reversal leaning is measured by the number of trials the participant took to learn the contingencies once they had realised the deck probabilities had reversed. Additionally, the post-reversal learning period included a measure of regressive errors. The number of regressive errors was defined as the number of times the previously favoured deck was selected during the post-reversal learning period (i.e. after the perseverative period had ended).

#### Temporal Difference Reinforcement Learning Model

We modelled participants’ choice behaviour as a function of their previous choices and rewards using a temporal difference reinforcement learning algorithm (Sutton and Barto, 1998). This allows us to track trial-and-error learning for each participant, during each task stage, in terms of a subjective expected value for each deck. On each trial *t*, the probability that deck *c* was chosen was given by a soft-max probability distribution,

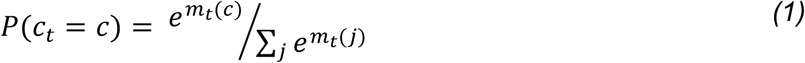

where *m_t_*(*c*) is the preference for the chosen deck and *j* indexes the four possible decks. The preference for the chosen deck was comprised of the participant’s expected value of that deck on that trial, *V_t_*(*c*), multiplied by the participant’s individual value impact parameter β (equivalent to

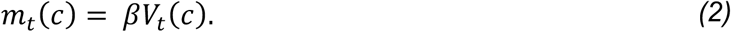

The parameter β describes the extent to which trial-by-trial choices follow the distribution of the expected values of the decks: a low β indicates choices are not strongly modulated by expected value, being effectively random with respect to this quantity (i.e. participants are not choosing based exclusively on value, and are effectively exploring all options); conversely, a high β indicates choices largely follow expected value (i.e. participants choose the deck with the highest expected value; exploitation).

To update the subjective value of each deck, a prediction error was generated on each trial, pet based on whether participants experienced a reward or a loss (*reward_t_*, = +1 or −1 respectively). The expected value of the chosen deck was subtracted from the actual trial reward to give the prediction error:

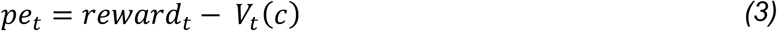

Studies have shown that individuals differ in the degree to which they learn from better than expected outcomes (positive prediction errors) and worse than expected outcomes (negative prediction errors) (Gray, 1970; Niv et al., 2012; Christakou et al., 2013; Bull et al., 2015). To account for this, two learning rate parameters were used to model sensitivity to prediction errors in updating the expected values: the weight of learning from better than expected outcomes (learning rate from positive prediction errors: *η*^+^) and the weight of learning from worse than expected outcomes (learning rate from negative prediction errors: *η*^−^). For example, individuals who are reward seeking will place a high weight on the former, whereas those who are loss-aversive will place a high weight on the latter. The prediction error on each trial was multiplied by either the positive (*η*^+^) or negative (*η*^−^) learning rate and used to update the value of the chosen deck.

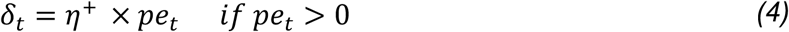

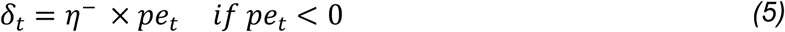

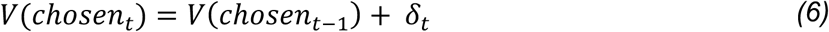

Thus, the model has three parameters of interest (β, *η*^+^ and *η*^−^). In psychological terms, β captures the degree to which the subjective value of the chosen deck influenced decisions, while the learning rates capture the individual’s preference for learning from positive (*η*^+^) or negative (*η*^−^) prediction errors to guide choice behaviour during this task.

#### Model Fitting

The model was fit per participant to provide parameters that maximised the likelihood of the observed choices given the model (individual maximum likelihood fit; Daw, 2011). The reward value was updated as 1 (win) or −1 (loss). Subjective value was initialised at zero for all decks and the initial parameter values were randomised. To ensure the model produced consistent, interpretable parameter estimates, *η* was limited to between 0 and 1 and β was assumed positive. The parameters were constrained by the following distributions based on Christakou et al (2013):

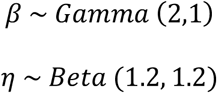

The model was fit separately over the trials encompassing round 1 (R1, initial learning) and round 2 (R2, perseverative trials and post-reversal learning, denoted as reversal learning). This was done to capture the change in influence of the model parameters from initial learning to reversal learning. The model was not fit over the perseverative trials separately as the average number of perseverative trials was too small to generate a stable model fit.

Traditionally, to investigate the fit of a temporal difference reinforcement learning model the Bayesian information criterion (BIC) is used. The BIC is a post hoc fit criterion which looks at the adequacy of a model whilst penalising the number of parameters used. A lower number indicates a better fit (Steingroever et al., 2016). However, the BIC is generally used to compare different models, rather than model fits over different sets of data, and will penalise different sized data sets. Alternatively, the corrected likelihood per trial (CLPT) can be used. The CLPT is a more intuitive measure of fit that takes into account the number of trials completed without penalising different sized data sets. The CLPT varies between 0 and 1, with higher values indicating a better fit (Leong and Niv, 2013; Niv et al., 2015). Wilcoxon signed-rank tests showed there was no significant difference between the CLPT values for the model fit over round 1 (Mdn = 0.23) and round 2 (Mdn = 0.23; *Z*= −1.308, p = 0.191). Additionally, there was no significant difference between the BIC values for the model fit over round 1 (M = 75.7, SD = 45.5) and round 2 (M = 90.9, SD = 43.6; *t*(33) = −1.533, p = 0.135, r = 0.26).

To summarise, the model fit equally well across rounds. Therefore, differences in parameter estimates across the task can be examined.

### Magnetic Resonance Spectroscopy

#### Data Acquisition

Data was collected at the University of Reading on a Siemens Trio 3T MRI scanner using a transmit-receive head coil. A high-resolution whole-brain T1 structural image was acquired for voxel placement using an MPRAGE sequence parallel to the anterior-posterior commissure line (176 × 1mm slices; TR = 2020ms; TE = 2.9ms; FOV = 250mm).

Voxels were placed in either the left or right dorsal striatum (DS), ventral striatum (VS) and the cerebellum, with hemisphere placement and order of measurements counterbalanced across participants. Anatomy was used to guide voxel positioning. The top of the DS was identified by slice-by-slice examination of the structural scan. The slice below the slice where the top of the striatum was no longer visible was selected and the top of the voxel was aligned with this slice. The same technique was applied for the VS voxel. The slice above the slice where the bottom of the striatum could no longer be seen was selected and used for alignment of the VS voxel. The cerebellum voxel was placed as high in the superior cerebellar vermis as possible whilst ensuring only cerebellar tissue was contained in the voxel. The superior cerebellar vermis was chosen as it has been shown to have the lowest variability in both inter and intra subject metabolite ratios as measured with ^1^H-MRS at rest (Currie et al., 2013). All voxels were visually inspected to ensure minimal cerebrospinal fluid was included in the voxels.

A PRESS sequence was used to acquire data from the three separate voxel positions (voxel size = 10mm x 15mm x 15mm; TR = 2000ms; TE = 30ms). 128 spectra were collected and averaged for each area. A water-unsuppressed spectrum was also obtained from each area for data processing, which consisted of an average of 15 spectra. Spectra were obtained for all participants from the cerebellum and the DS. The spectrum obtained from the VS was only usable from 34 participants due to noise levels. The SIEMENS Auto Align Scout was used in between each scan to adjust the voxel position based on the actual head position of the participant. This was used to correct for participant motion and minimize the variability of the voxel position.

**Figure 3:**
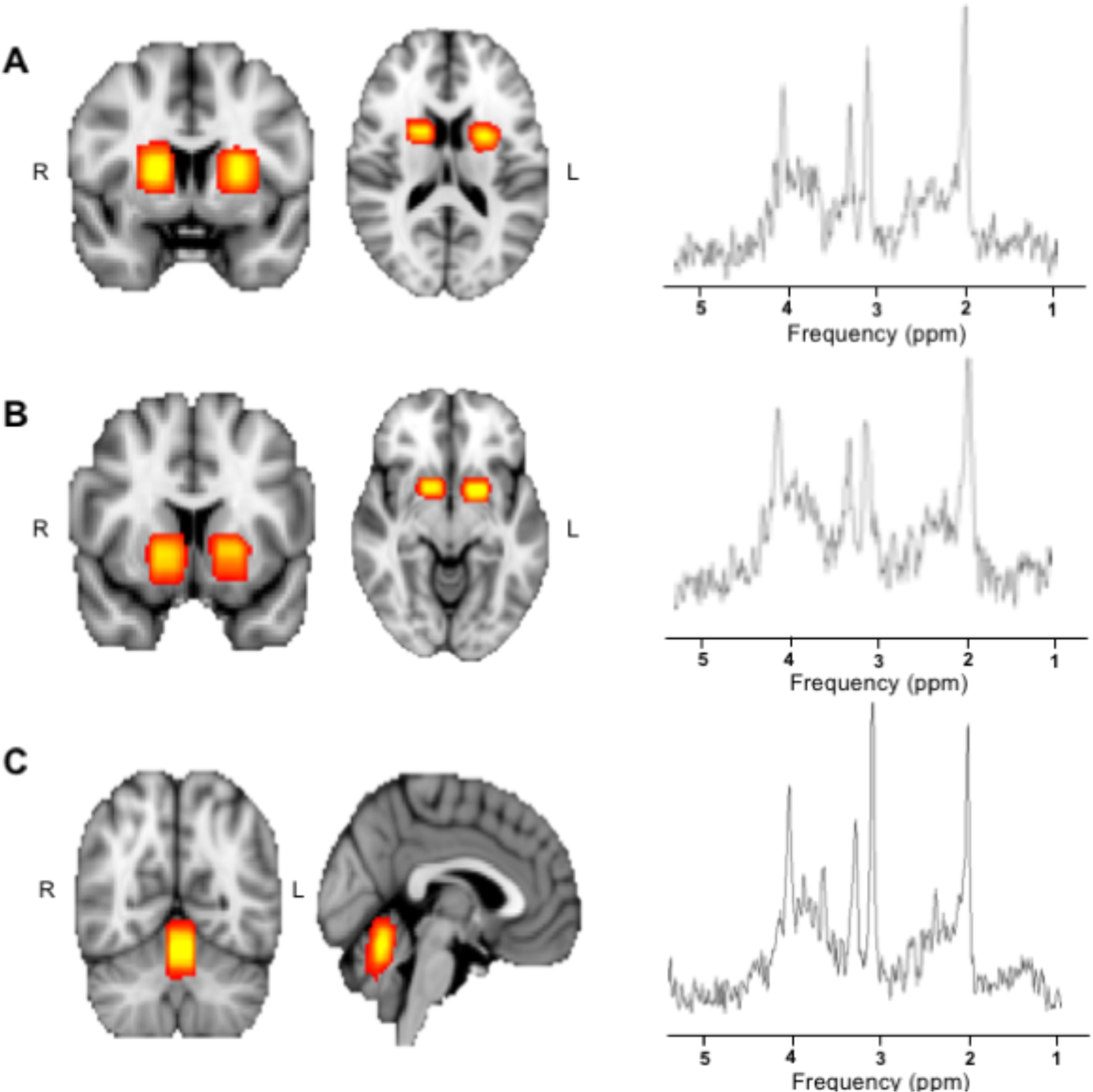
Location of voxels and example spectra. Heat maps showing the sum of the ^1^H-MRS voxels over all subjects in MNI space, along with a representative spectrum from a single subject (A = Dorsal Striatum, MNI coordinates: −3.41, 2.37, 11.16; B = Ventral Striatum, MNI coordinates: −2.99, 5.92, −3.93; C = Cerebellum, MNI coordinates: −2.10, −61.03, 19.20).

#### Structural Segmentation

Structural scans were processed using FSL version 5.0.8 (Smith et al., 2004; Jenkinson et al., 2012). First, the skull was removed using the brain extraction tool (BET) (Smith, 2002). Images were segmented into three separate tissue types: grey matter (GM), white matter (WM) and cerebrospinal fluid (CSF) using the FAST tool (Zhang et al., 2001). The coordinates and dimensions of the voxel were then superimposed on these images and the proportion of each of the three tissue types contained within the voxel was calculated.

#### Quantitation

Data was processed in the time domain using Java-Based Magnetic Resonance User Interface (jMRUI software version 5.0 (http://www.mrui.uab.es/mrui); Naressi et al., 2001). Phase correction was performed using the corresponding water spectrum from each area. Each spectrum was then apodized using a Gaussian filter of 3Hz to improve signal quality, reduce noise and reduce effects of signal truncation (Jiru, 2008). The residual water peak was removed using the Hankel-Lanczos Singular Value Decomposition (HLSVD) filter tool.

Metabolite models were generated using the software Versatile Simulation, Pulses and Analysis (VEsPA (https://scion.duhs.duke.edu/vespa/project); Soher, Semanchuk, Todd, Steinberg, & Young., 2010). 14 typical brain metabolites (Acetate, Aspartate, CHO, Creatine, Gamma-Aminobutyric Acid (GABA), Glucose, Glutamate, Glutamine, Lactate, Myo-inositol, N-acetyl Aspartate (NAA), Phosphocreatine, PC & GPC, Scyllo-inositol, Succinate, Taurine) were simulated at a field strength of 3T using a PRESS pulse sequence (TE1 = 20ms, TE2 = 10ms, main field = 123.25MHz). For initial analyses, CHO was modelled separately from PC+GPC based on the method described in Bell et al., 2018. Additionally, the sum of the three peaks (total choline, tCHO) was included in the analyses for comparison. If tCHO produced similar results to CHO, it would potentially suggest that there may not be a need to separate the three peaks, or that the quantitation method is not separating CHO effectively.

The jMRUI tool Accurate Quantification of Short Echo time domain Signals (AQSES) was used for automatic quantification of spectra signals. AQSES was applied using the method described in Minati, Aquino, Bruzzone, & Erbetta, 2010. To correct for any chemical shift displacement, the spectrum was shifted so that the peak for n-acetyl-aspartate (NAA) was at 2.02ppm. The frequency range selected for processing was limited to 0-8.6ppm (equal phase for all metabolites, begin time fixed, delta damping (−10 to 25Hz), delta frequency (−5 to 5Hz), no background handling, 0 truncated points, 2048 points in AQSES and normalisation on). Based on common practice in the field, values with a CRB higher than 30% were excluded on a case by case basis.

Metabolite concentrations were calculated for CHO, PC+GPC, tCHO, NAA and total creatine (tCR, creatine + phosphocreatine), correcting for partial-volume and relaxation effects, using the formula described in Gasparovic et al., 2006).

### Experimental Design and Statistical Analysis

Statistical analysis was performed using SPSS (IBM Corp. Released 2013. IBM SPSS Statistics for Windows, Version 22.0. Armonk, NY: IBM Corp).

The relationships between model parameters and behaviour, along with model parameters and metabolite levels and behaviour and metabolite levels was assessed using correlation analysis. The distribution of the data was analysed using measures of skewness and kurtosis, along with the Shapiro-Wilk test. When the assumptions of normality and homogeneity were met, Pearson’s correlation (*r*) was used to assess correlations. When assumptions of normality were not met, Kendall’s Tau (*r*□) was used to assess correlations, as it provides a better estimation of the correlation in a small sample size compared to other non-parametric methods (Field, 2009). Where appropriate, hierarchical multiple regression analysis was used to assess the variance explained by metabolite levels in behaviour, after the model parameters were accounted for.

#### Confounding Variables

There were no significant differences in metabolite levels between hemispheres, therefore the results were combined across hemisphere of acquisition.

To examine if variations in the metabolite values might be caused by differing proportions of tissue composition, correlations were performed between CCC levels and proportion of grey and white matter present in the voxel. Additionally, metabolite values were checked against the water signal for the same reason. No significant correlations were found between CCCs and grey/white matter content, indicating any variance seen is generated by differing metabolite levels. The water signal significantly correlated with DS tCHO (*r*□ (35) = −0.348, p = 0.003) and VS PC+GPC (*r*□ (32) = − 0.270, p = 0.001). Therefore, analyses involving DS tCHO or VS PC+GPC were corrected for this source of variance using partial correlations. No other significant correlations were seen between the water signal and metabolite levels of interest.

There is evidence that metabolite levels in the brain can vary based on time of day (Soreni et al., 2006) and age (Pfefferbaum et al., 1999; Reyngoudt et al., 2012). Therefore, all metabolites were checked against these two variables to ensure this was not a source of variance. Time of day significantly correlated with DS tCHO (*r*□ (35) = 0.249, p = 0.038) and cerebellum tCHO (*r*□ (31) = 0.285, p = 0.026). Therefore, analyses involving DS tCHO or cerebellum tCHO were corrected for this source of variance using partial correlations. No other significant correlations were seen between metabolite levels and time of day or age of participant.

#### Controls

The cerebellum was used as a control to demonstrate the regional specificity of results. None of the effects were present in the cerebellum and therefore these results are not reported further. NAA and tCR were used as controls to demonstrate the neurochemical specificity of the results (i.e. that the relevant individual differences were specific to choline and not to spectrum-wide inter-individual differences). None of the effects were present in either NAA or tCR and therefore these results are not reported further. Furthermore, none of the reported effects were found when using tCHO as a measure of cholinergic availability and therefore these results are not reported further.

## RESULTS

### Behavioural Results

Twenty-two (22) participants reached criterion during both rounds (i.e. they reached criterion both during initial learning and after the reversal) and were included in the analysis.

#### Model parameters and performance

A reinforcement-learning model was used to disentangle components of learning that contribute to overall behaviour. We looked at three parameters of interest, the learning rates from positive (*η*^+^) and negative (*η*^−^) prediction errors, and the overall impact of subjective value of the deck on the participants choice (value impact parameter, β). To test the contribution of the model parameters to behaviour, we looked at correlations between behaviour (as measured by trials to criterion, number of perseverative trials and number of regressive errors) and the corresponding model parameters, i.e. behaviour during initial learning was correlated with model parameters fit over the initial learning period, and likewise for the reversal learning period.

**Table 1:** Performance variables

|  | *Average Number of Trials* | *SD* |
| --- | --- | --- |
| **Initial Learning** | 44 | 28 |
| **Reversal Learning** |  |  |
| **Perseveration Period** | 12 | 8 |
| **Post Reversal Learning** | 35 | 22 |
| **Regressive Errors** | 7 | 6 |
| **Total** | 47 |  |

**Table 2:** Estimates of model parameters

| | $\eta^+$ | $\eta^-$ | $\beta$ |
| --- | --- | --- | --- |
| **Initial Learning** | 0.37<br>(SD = 0.30) | 0.42<br>(SD = 0.31) | 1.44<br>(SD = 0.56) |
| **Reversal Learning** | 0.24<br>(SD = 0.35) | 0.31<br>(SD = 0.27) | 1.37<br>(SD = 0.97) |

Note: $\eta^+$ = learning rate from positive prediction errors; $\eta^-$ = learning rate from negative prediction errors; $\beta$ = impact of subjective value on choice.

Table 3 shows the correlation coefficients for the relationships between model parameters and behaviour. Faster initial learning (low number of trials to criterion) was associated with a higher learning rate from positive prediction errors (*r*(22) = −0.439, p = 0.041) and a higher value impact parameter (*r*(21) = −0.536, p = 0.012). A lower number of perseverative trials was associated with a higher learning rate from negative prediction errors (*r*□(22) = −0.527, p = 0.012). As was the case during initial learning, during post-reversal learning (after the reversal has been identified) a lower number of trials taken to reach criterion was associated with a higher learning rate from positive prediction errors (*r*□ (22) = −0.335, p = 0.03), and a higher value impact parameter (*r*□ (22) = − 0.352, p = 0.022). Additionally, during post-reversal learning, a lower number of regressive errors was associated with a higher learning rate from positive prediction errors (*r*□ (22) = −0.355, p = 0.023) and a higher value impact parameter (*r*□□ (22) = −0.337, p = 0.031).

**Table 3:** Correlation coefficients for relationships between model parameters and behaviour

| | $\eta^+$ | $\eta^-$ | $\beta$ |
| --- | --- | --- | --- |
| **Initial Learning (TTC)** | -0.439* | -0.218 | -0.536* |
| **Reversal Learning** |  |  |  |
| **Perseverative Errors** | -0.176 | -0.527* | 0.132 |
| **Post Reversal Learning (TTC)** | -0.335* | 0.322 | -0.352* |
| **Regressive Errors** | -0.355* | 0.292 | -0.337* |

Note: $\eta^+$ = learning rate from positive prediction errors; $\eta^-$ = learning rate from negative prediction errors; $\beta$ = value impact parameter; \* $p < 0.05$ .

#### Effects of trait impulsivity on performance

To investigate the influence of impulsivity on decision making, we looked at correlations between impulsivity (total BIS-11 score) and measures of behaviour (including model parameters) in learners. Higher impulsivity levels were associated with a lower number of perseverative errors (*r*(22) = −0.470, p = 0.027). No other measures of behaviour correlated with impulsivity.

#### Summary

Faster initial learning was indexed by both higher learning rates from positive prediction errors (R1*η*^+^) and higher value impact parameters (R1β). Reduced numbers of perseverative trials were associated with higher learning rates from negative prediction errors (R2*η*^−^) and higher impulsivity levels. Similar to initial learning, faster post-reversal learning was associated with higher learning rates from positive prediction errors (R2*η*) and higher value impact parameters (R2β). Additionally, during post-reversal learning, lower numbers of regressive errors were associated with higher learning rates from positive prediction errors (R2*η*^+^) and higher value impact parameters (R2β).

### Spectroscopy Results

One participant was excluded from spectroscopy analysis due to issues with segmentation of the structural scan.

**Table 4:** Average metabolite levels in the DS

|  | *CHO* | *PC+GPC* | *tCHO* | *NAA* | *tCR* |
| --- | --- | --- | --- | --- | --- |
| **Learners** | 0.15<br>(SD = 0.20) | 0.27<br>(SD = 0.10) | 0.42<br>(SD = 0.12) | 8.73<br>(SD = 0.77) | 11.58<br>(SD = 1.74) |
| **Non-Learners** | 0.11<br>(SD = 0.16) | 0.36<br>(SD = 0.14) | 0.46<br>(SD = 0.10) | 8.83<br>(SD = 2.37) | 11.80<br>(SD = 2.31) |

Note: CHO = choline, PC+GPC = phosphocholine and glycerophosphocholine, tCHO = total choline, NAA = n-acetyl aspartate, tCR = total creatine.

**Table 5:** Average metabolite levels in the VS

|  | *CHO* | *PC+GPC* | *tCHO* | *NAA* | *tCR* |
| --- | --- | --- | --- | --- | --- |
| **Learners** | 0.24<br>(SD = 0.17) | 0.27<br>(SD = 0.12) | 0.5<br>(SD = 0.17) | 5.39<br>(SD = 1.97) | 12.02<br>(SD = 2.26) |
| **Non-Learners** | 0.23<br>(SD = 0.17) | 0.25<br>(SD = 0.14) | 0.48<br>(SD = 0.16) | 5.45<br>(SD = 1.54) | 11.13<br>(SD = 3.95) |

Note: CHO = choline, PC+GPC = phosphocholine and glycerophosphocholine, tCHO = total choline, NAA = n-acetyl aspartate, tCR = total creatine.

#### Group Comparisons

To investigate whether average levels of CHO in the striatum relate to task performance, the average levels were compared between learners and non-learners. There was no significant difference in CHO levels between learners and non-learners in either the DS or the VS.

#### Correlations with Behaviour

To further investigate the relationship between average metabolite levels and task performance, correlations were performed between metabolite levels and measures of performance in learners (numbers of trials to criterion and model parameters).

##### Initial Learning

No significant correlations were seen with measures of performance in round 1 (trials to criterion, R1*η*^+^ or R1β) and average levels of CHO in the DS.

VS CHO did not correlate with trials to criterion in round 1. However, low levels of CHO in the VS were associated with higher learning rates from positive prediction errors (*r*(20) = −0.625, p = 0.003; Figure 4A) and lower value impact parameters (*r*□(19) = 0.555, p = 0.014; Figure ***4***B).

**Figure 4:**
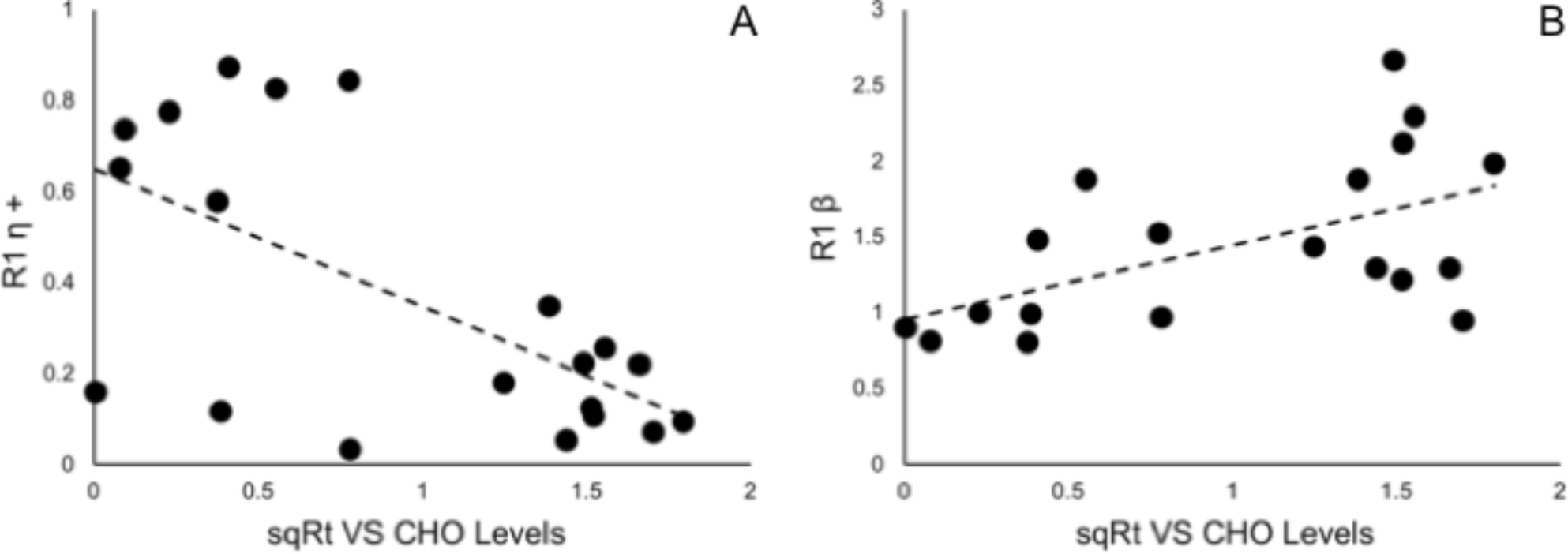
Correlations between VS CHO levels and performance during initial learning **A:** Negative correlation between learning rate based on positive prediction errors derived from round 1 (R1*η*+) and levels of CHO in the VS (r(20) = −0.625, p = 0.003). **B:** Negative correlation between impact of participant’s subjective value on their future choice derived from round 1 (R1β) and levels of CHO) in the VS (r(19) = 0.555, p = 0.014). VS: Ventral Striatum; CHO: Choline.

##### Perseverative Trials

A lower number of perseverative trials was associated with lower levels of DS CHO (*r*□ (21) = 0.367, p = 0.021; Figure 5A). The opposite effect was seen with DS PC+GPC (r(21) = −0.447, p = 0.042. Additionally, higher learning rates from negative prediction errors were associated with lower DS CHO levels (*r*□ (21) = −0.371, p = 0.019; Figure 5B). This result is specific to DS CHO, with no other DS metabolites found to correlate with learning rates from negative prediction errors. Additionally, VS CHO was not found to correlate with either the number of perseverative trials or learning rates from negative prediction errors.

**Figure 5:**
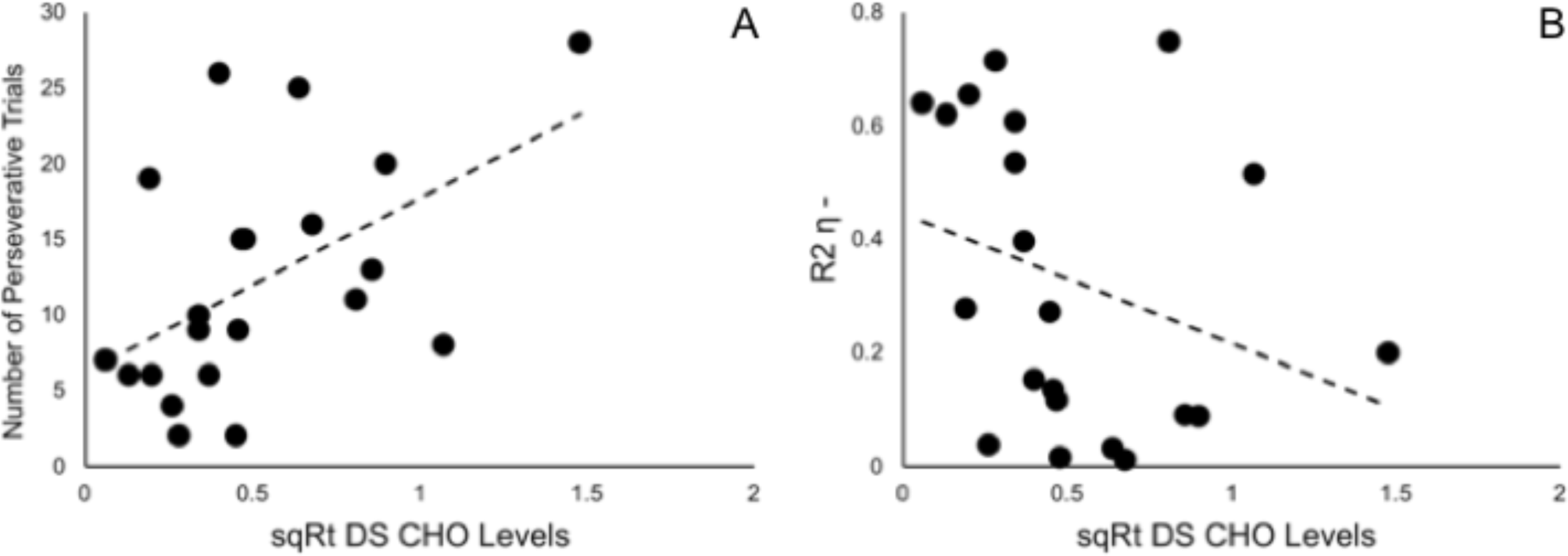
Correlations between DS CHO levels and performance during reversal **A:** Positive correlation between the number of perseverative trials and levels of CHO in the DS (rD (21) = 0.367, p = 0.021). **B:** Negative correlation between the learning rate based on negative prediction errors derived from round 2 (R2*η*-) and levels of CHO in the DS (*r*□ (21) = −0.371, p = 0.019). DS: Dorsal Striatum; CHO: Choline.

After establishing an association between CHO levels and reversal performance, we wanted to examine whether CHO contributed to reversal efficiency over and above behavioural and personality variables. Using a hierarchical multiple regression, we first modelled the contribution of variance from learning rates from negative prediction errors and total BIS scores to the variance in the number of perseverative trials (Model 1; F(2,18) = 9.460 p = 0.002, R^2^ = 0.512; Table 6). The second model looked at whether the addition of DS CHO would explain significantly more variance, over and above that explained by learning rates from negative prediction errors and total BIS score (Model 2; F(3,17) = 9.574 p = 0.001, R^2^ = 0.628; Table 6).

**Table 6:**
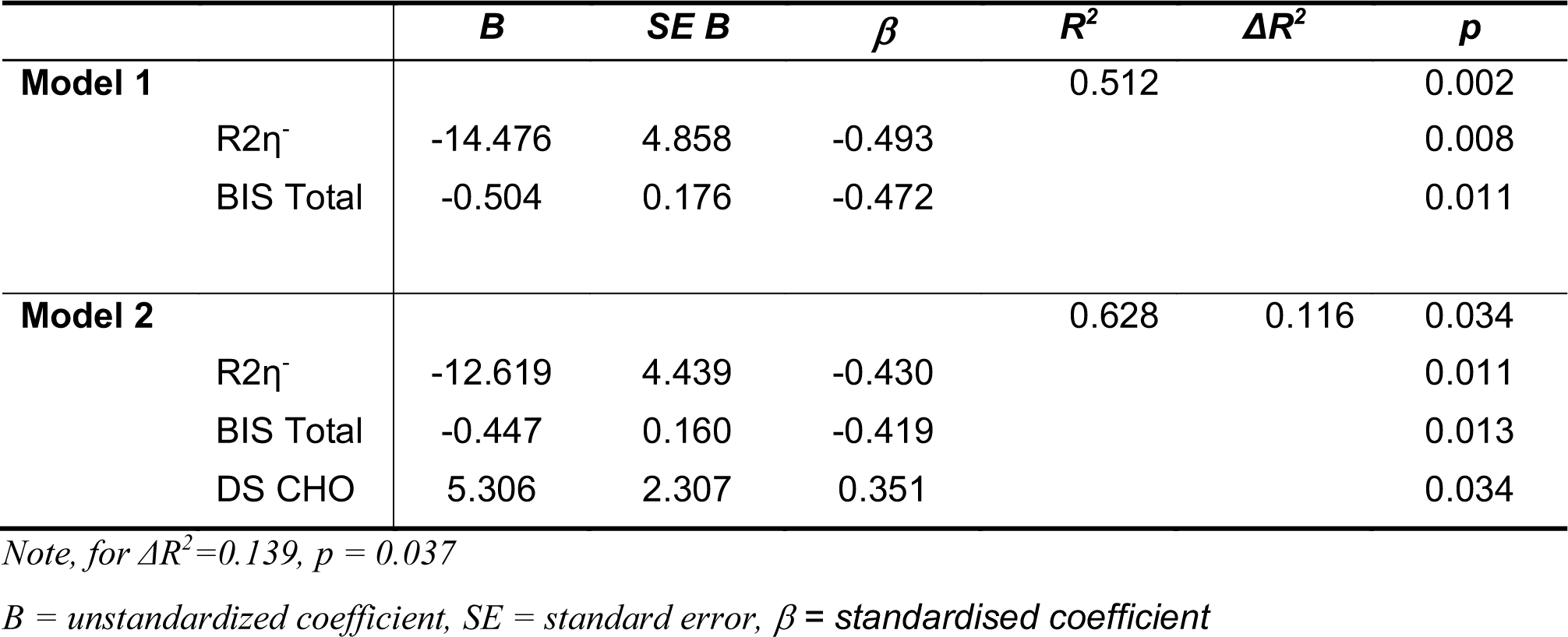
Summary of hierarchical regression analyses for variables predicting perseveration

The amount of variance in the number of perseverative trials explained by learning rates from negative prediction errors was significant in both Model 1 (β = −0.493, t(18) = −2.980, p = 0.008; Table 6) and Model 2 (β = −0.430, t(17) = −2.843, p = 0.011; Table 6). Additionally, total BIS score also explained a significant amount of variance in both Model 1 (β = −0.472, t(18) = −2.855, p = 0.011; Table 6) and Model 2 (β = −0.419, t(17) = −2.787, p = 0.013; Table 6).

In Model 2, DS CHO also explained a significant amount of variance in the number of perseverative trials (β = 0.351, t(17) = 2.300, p = 0.034; Table 6). The addition of DS CHO to the model increased R^2^ by 0.116 and this increase was statistically significant (F(1,23) = 5.291, p = 0.034; Table 6).

To assess the specificity of this result, DS PC+GPC was also included in the model. However, analysis of multicollinearity diagnostics showed a tolerance of 0.175, which is below the acceptable value of 0.2. This is due to the strong significant correlation between DS CHO and DS PC+GPC (*r*□ (21) = −0.667 p < 0.001). As a result, including the two variables in the same regression model would violate the assumption of multicollinearity and the regression model would not be able to provide unique estimates of the regression coefficients, as each will account for overlapping variance (Field, 2009). Therefore, we instead repeated the hierarchical regression with DS PC+GPC in place of DS CHO. The amount of variance explained by DS PC+GPC was not significant (β = − 0.301, t(17) = −1.900, p = 0.075). The addition of DS PC+GPC to the model increased R^2^ by 0.085 and this increase was not statistically significant (F(1,23) = 3.611, p = 0.075). This indicates that DS CHO levels can explain part of the variance in the number of perseverative trials, however DS PC+GPC levels cannot.

##### Post Reversal Learning

No significant correlations were seen with either DS or VS CHO levels and measures of performance during post reversal learning (trials to criterion, R2*η*^+^ or R2β). Additionally, there were no significant correlations between DS or VS CHO levels and the number of regressive errors.

#### Summary

In the DS, average CHO levels were associated with performance during reversal, but not during initial learning. There was a significant positive correlation between DS CHO levels and the number of perseverative trials, and a significant negative correlation between DS CHO levels and learning rates from negative prediction errors (R2*η*^−^). Additionally, DS CHO levels explained variance in the number of perseverative trials over and above that explained by learning rates from negative prediction errors.

In the VS, average CHO levels were not associated with performance during reversal learning. Although VS CHO levels were not associated with the speed of initial learning, there was a significant positive correlation between VS CHO levels and learning rates from positive prediction errors, and a significant negative correlation between VS CHO levels and the value impact parameter during initial learning.

## DISCUSSION

We used ^1^H-MRS to investigate the relationship between average CHO levels in the human striatum at rest and performance during probabilistic reversal learning. Here we show that baseline levels of CHO in the human DS are associated specifically with individual differences in reversal learning efficiency, but not in initial learning, and that this effect is specific to the dorsal, but not the ventral striatum.

Behaviourally, we show that faster initial learning is indexed by a higher learning rate from positive prediction errors (*η*^+^) and a higher value impact parameter (β). Therefore, during this period, participants are using wins and expected value to guide their choices. This is also seen during the post-reversal learning period, in which faster post-reversal learning is indexed by higher learning rates from positive prediction errors (*η*^+^) and higher value impact parameters (β). Faster reversal (less perseveration), however, was indexed by higher learning rates from negative prediction errors (*η*^−^) only. During this period, participants must now pay increased attention to worse than expected outcomes in o*r*□er to identify the change in contingencies. Therefore, to adapt to changes in task structure, participants adapt their strategy by altering the weight of learning from prediction errors based on rewa*r*□ history.

The learning rate for negative prediction errors, accounting for trait impulsivity, explained a significant amount of variance during perseveration, providing a simple mechanism to explain reversal efficiency. However, average DS CHO levels explained variance in the number of perseverative trials over and above this original model. This suggests a more complex mechanism in which perseveration is influenced, in part, by the learningrate from negative prediction errors (which can change due to task demand) and by resting levels of DS CHO. Indeed, Franklin & Frank, 2015 showed that a model which takes into account cholinergic activity performs better on a reversal learning task than a model based solely on dopamine prediction error signalling.

Our results indicate that participants who were quicker to reverse had lower average levels of DS CHO, suggesting that low trait levels of DS CHO are beneficial for reversal learning. Based on evidence that ACh efflux increases during reversal learning (Ragozzino et al., 2009; Brown et al., 2010), this suggests two potential mechanisms. Firstly, lower levels of DS CHO at rest could reflect lower levels of ACh at rest. This is also supported by evidence from the animal literature, which has shown a positive correlation between ACh levels at rest as measured by microdialysis and average CCCs as measured by ^1^H-MRS (Wang et al., 2008). Additionally, higher levels of CHO availability have been shown to lead to higher levels of ACh release, implying a positive correlation between the two metabolites (Koshimura et al., 1990). Based on this notion, the findings here suggest that lower levels of ACh at rest may be beneficial for reversal learning because they enable a higher contrast between ACh levels at rest and during reversal learning. However, it is important to note that Wang et al. (2008) modelled all three CCCs as a single peak. It is likely that the relationship between CHO levels as measured by spectroscopy and ACh levels in the brain is not straightforward, and this interpretation should be considered with caution. Indeed, animal studies have shown the relationship between CHO and ACh can change based on neuronal firing and ACh requirement (Löffelholz, 1998; Klein et al., 2002). Furthermore, we have previously demonstrated a drop in CHO levels in the human DS during reversal learning, thought to reflect the sustained increase in ACh release seen in animal studies (e.g. Ragozzino et al., 2009). This drop is thought to be due to an increase in translocation of CHO uptake receptors in response to sustained neural firing (Bell et al., 2018). Though we have described the measurements in this study as “at rest”, cholinergic interneurons are tonically active, and therefore the relationship between CHO and ACh levels in the striatum will likely reflect a more complex dynamical relationship between the two.

The second potential mechanism supported by our findings is that lower levels of DS CHO at rest may result from a more efficient CHO uptake system. Mice carrying mutations in the gene coding for CHO uptake transporters have reduced neuronal capacity to both clear CHO and release ACh.

Moreover, performance on an attention task was impaired in these mice (Parikh et al., 2013). Additionally, in a study of frontal cortex cholinergic modulation during attention, humans with a gene polymorphism which reduces CHO transport capacity showed reduced activation in the prefrontal cortex during an attentional task. Furthermore, the pattern of activation predicted CHO genotype (Berry et al., 2015). Further work is needed to determine the relationship between CHO uptake, ACh release and reversal learning.

Disruption of cholinergic signalling in rodents typically results in an increase in regressive errors (Brown et al., 2010; Bradfield et al., 2013). However, here we found no association between DS CHO levels and the number of regressive errors. In humans, measures of individual differences in perseverative and regressive errors are likely to be confounded by individual differences in representation of the task structure. Rather than making perseverative and regressive errors based solely on feedback, the ability to flexibly alter response depends in part on a higher level representation of the task, which is thought to be maintained in frontal areas of the cortex (Armbruster et al., 2012). It should be noted that the basal ganglia-thalamo-cortical system has been shown to be modulated by the maintenance of task rules, with those with stronger representation of the task structure showing higher activation in the caudate and thalamus during a behaviour switch (Ueltzhöffer et al., 2015), indicating that representation of task structure likely modulates DS activity in response to the need for behavioural flexibility. Inevitably, caution is needed when translating evidence from rodent studies of learning to human studies. This emphasises the need to further develop non-invasive techniques for studying human neurochemistry *in vivo*

As predicted, and in line with evidence from the animal literature (Ragozzino et al., 2009), levels of CHO in the VS were not associated with reversal learning. However, VS CHO levels were associated with model parameters which contributed to initial learning. Though Ragozzino et al. demonstrated that ACh levels in the VS did not change during reversal learning, they did not test if they changed during initial learning. Successful learning requires the ability to learn from feedback, which is signalled by dopaminergic prediction error signalling in the VS (Schultz et al., 1997). The rodent VS has a higher density of cholinergic interneurons than the DS (Matamales et al., 2016) and changes in cholinergic activity are time locked to changes in dopaminergic activity, which is thought to enhance the contrast of prediction error signalling (Aosaki et al., 2010). Indeed, cholinergic activity in the VS has been linked with effective learning of a stimulus-outcome association (Brown et al., 2012), therefore, it is likely that cholinergic activity in the VS is involved in some aspect with goal-directed learning, and further studies should explore this contribution.

We used several controls to demonstrate that these effects are specific to CHO levels in the striatum, not least because our MRS application method is novel. We acquired data from a voxel in the cerebellum, geometrically identical to the striatal voxels. No learning effects were present in the cerebellum, demonstrating that our findings are specific to the striatum. Additionally, we also quantified two control metabolites (NAA and tCR) to ensure that the results were specific to the metabolite of interest, rather than a general measurement or region effect. None of the effects were seen in levels of NAA and tCR in the DS or VS. Importantly, none of the effects were seen when modelling all three peaks together (tCHO), highlighting once more the importance of separating CHO when using ^1^H-MRS to investigate individual differences in CCC levels.

In conclusion, ^1^H-MRS was used to demonstrate that average levels of CHO in the human DS are associated with performance during probabilistic reversal, but not during initial learning. This is in line with evidence from the animal literature and out own prior work with humans which suggests a specific role for cholinergic activity in the DS during reversal learning. These results provide evidence for the role of the human cholinergic striatum in reversal learning and behavioural flexibility more generally. Additionally, these findings further support the idea of using CHO levels as measured by ^1^H-MRS as a tool for non-invasive *in vivo* monitoring of both healthy human neurochemistry, as well as diso*r*□ers of the human cholinergic system.

## Acknowledgments

This study was supported by a Human Frontier Science Program (HFSP) grant (RGP0048/2012), and an Engineering and Physical Sciences Research Council (EPSRC) doctoral training grant (EP/L505043/1). The funders had no involvement in study design, in the collection, analysis, and interpretation of data, in writing the report, or in the decision to submit the paper for publication. The authors would like to thank Rosie Gillespie and Emma Davis for assistance with data collection.

